# Seasonal genotypic and phenotypic differentiation of a cosmopolitan freshwater diatom

**DOI:** 10.1101/2024.03.20.585933

**Authors:** Domiziana Cristini, Joseph B. Kelly, Pia Mahler, Lutz Becks

## Abstract

Most ecosystems are characterized by seasonality, which, through biotic and abiotic changes, influences species biomass dynamics. Recent studies have shown that ecologically important traits can evolve rapidly in response to environmental changes, resulting in eco-evolutionary dynamics with consequences for population and community dynamics. Evidence for seasonal effects on intraspecific variation is still scarce and understanding eco-evolutionary dynamics in the presence of seasonal fluctuations remains a major challenge. Following the phytoplankton spring bloom in Lake Constance, we investigated how seasonal changes influence the intraspecific diversity of *Asterionella formosa* both at genotypic and phenotypic levels. We found a high degree of genetic and phenotypic differentiation characterizing *Asterionella* population, explained by a clustering of the isolates into early and late spring according to lake thermal stratification. Yet, most traits related to environmental parameters as well as fitness in different seasonal environments did not show a clear response to seasonality (i.e., temperature and nutrients), indicating that seasonal changes in biotic interactions (i.e., parasitic chytrids) were the major drivers of the observed seasonal shift in *Asterionella* genotypes. Our results highlight the importance of studying eco-evolutionary processes for understanding variations in population and community dynamics in response to seasonal environmental fluctuations.

## Introduction

Seasonality can be an important driver of species abundance and distribution and has been observed in terrestrial [e.g., 1,2] and aquatic ecosystems [e.g., 3,4]. Periodic changes in the biotic and abiotic environment can often be linked to changes in species biomasses and functional groups. For example, in temperate lakes the biomass of phytoplankton and zooplankton has been described to follow the seasonal pattern defined in the PEG model (Plankton Ecology Group model [5]). The model predicts a spring phytoplankton bloom led by small and highly competitive species under high nutrient concentrations and water mixing dynamics, followed by a biomass minimum driven by zooplankton grazing. In summer, when nutrients are low and the water column is stratified, phytoplankton species that are larger and slower-growing but have a high nutrient affinity become abundant. Although these seasonal changes in plankton communities are well documented, observations often differ from the patterns predicted by the PEG and other models, for example for a lake over several years [6].

Studies of seasonal plankton dynamics typically focus on functional groups and species, but intraspecific variation has rarely been considered and are not integrated into the PEG model [6], even though it can have important consequences for communities [7–9]. Intraspecific variation can be relevant if shifts in genotype frequencies (= evolution) affect communities to a similar extent as ecology resulting in eco-evolutionary dynamics [10–13]. Examples for such dynamics in natural systems that are driven by annual seasonality are scarce. Schaffner et al. [14] reported the occurrence of rapid evolution in a *Daphnia* species during a single season in Lake Oneida. This seasonal adaptive evolution followed a change in food quality from spring to summer; an increase in the abundance of cyanobacteria, which selected for toxin-resistant clones and consequently altered the population dynamics of *Daphnia*. Similar genotype and population dynamics in phytoplankton natural communities with strong seasonal variation have not been well studied. Apart from this study, there is currently little data and understanding of how evolutionary and ecological processes drive the dynamics of phytoplankton communities in the presence of seasonality. Some studies have documented genetic differentiation of plankton populations across seasons (e.g. for diatoms: [15–18]; for *Daphnia*: [14,19]), but very few studies have linked the observed genetic diversity to traits and fitness consequences.

The observation of genetic and phenotypic variation within phytoplankton populations [12,20] and the strong seasonal patterns in temperate lakes (e.g. temperature, nutrient concentration, predation, parasitism) [5,6] suggest that intraspecific variation may give rise to eco-evolutionary dynamics and thus contribute to population dynamics. In the presence of eco-evolutionary dynamics, a largely untested expectation is that changes in genotypic and trait variation among individuals of the same species following seasonal changes will have consequences for fitness and thus may contribute to population growth. Here we tested this prediction by linking genotypic and phenotypic information from individual isolates of the cosmopolitan freshwater planktonic diatom *Asterionella formosa*, isolated from Lake Constance during the 2021 spring bloom. In spring the lake underwent a gradual increase of water temperature and light, with a simoultaneous decrease of nutrients and an increasing predation/parasitic pressure. During this period *A. formosa* is a dominant species. *A. formosa* reproduces mainly asexually [21] and we can assume clonal reproduction throughout the season, allowing us to isolate individual genotypes and propagating them clonally. We used single nucleotide polymorphisms (SNPs) obtained from whole-genome sequencing to assess the genetic diversity of 33 *Asterionella* isolates. For seven of these isolates, we measured phenotypic variation and estimated fitness under a range of temperature and nutrient conditions mimicking the important environmental conditions at the time of isolation (environmental temperature and nutrient conditions). The traits covered the physiological traits considered to be particularly susceptible to the main seasonal environmental stresses (i.e., nutrients, temperature, parasitic infection) [5]: maximum growth rate, carrying capacity, P-and Si half-saturation constants, sinking rate, cell length and temperature optimum.

## Methods

### Study site and sampling methodology

Lake Constance is a deep warm-monomictic peri-alpine lake situated at the border of three countries: Germany, Switzerland and Austria. The lake underwent re-oligotrophication from 1979 to mid 90s, and has since persisted in an oligotrophic state. During the spring bloom 2021 we sampled Upper Lake Constance weekly from the beginning of March (03.03.21) to the end of June (30.06.21) consistently in the same location (47°45’19.2“N 9°07’48.9”E). Temperature was measured from the surface to the bottom (0-100 m) using an RBR *maestro*³ sensor. On two sampling dates (15.05.2021 and 19.05.2021) temperature data were missing, and we used the values measured on the same day using a bbe Moldaenke FluoroProbe. We collected samples for assessing phytoplankton abundance and nutrients concentration using a messenger-released Free Flow Water Sampler 5 L (HYDRO-BIOS, Altenholz, Germany) over the uppermost water layer (0-20m, euphotic zone). Water samples for phytoplankton counting were immediately fixed with Lugol solution (0.62% final concentration). The same samples were also used to assess infection on *Asterionella formosa* by chytrids, which are fungal parasites known to be host specific and highly infective [22]. The counting of phytoplankton and chytrids was carried out under an inverted microscope following the standard Utermöhl protocol [23]. Water for the dissolved nutrients was filtered immediately upon collection using 0.45uM Whatman filters and stored in the dark until reaching the laboratory (∼2 hours). Nitrate (NO_3_) and phosphorus (P-PO_4_) were measured on the same day, while silicate (Si-SiO_2_) was stored at −20°C until measurement. Standard spectrophotometric methods (SpectroquantⓇProve 600, Merck) were applied to measure NO_3_ [24], P-PO_4_ [25] and Si-SiO_2_ [26]. Samples for silica measurements were stored and processed in plastic containers to avoid any contamination. Analytical blanks were analyzed prior to empirical measurements to correct for background contamination. All laboratory wares used for nutrient analysis were acid-washed prior to usage.

### Asterionella formosa *cultures*

We collected water samples for *Asterionella* isolation using a vertical plankton net-tow from 0 to 20 m. The samples were transported in the dark to the laboratory, where single colonies of *A. formosa* were immediately isolated under an inverted microscope and cultures established in WC medium [27]. We were able to successfully isolate and establish a total of 109 algal cultures over the entire sampling period. Due to experimental constraints and handling, 33 isolates were processed for molecular analysis from both early (03.03.21, 31.03.21 and 06.05.21) and late (09.06.21, 16.06.21, 23.06.21 and 30.06.21) spring and 7 from these cultures were randomly chosen (4 samples from 31.03.21, 3 samples from 30.06.21) and used to assess phenotypic variability. Isolates were maintained in non-axenic cultures at 14°C. Light intensity was set at 45 µmol m^−2^ s^−2^ (PAR) with a smooth light reduction to 10% every 20 min for 90 seconds to simulate environmental conditions. Cultures were refreshed every four weeks to assure no nutrient limitation.

### DNA extraction

100 mL of each target *Asterionella* culture was pelletized at 3260 x g for 20 min at 20°C. The pellet was transferred to 2.5-mL Eppendorf tubes and immediately frozen at −80°C until DNA extraction. DNA was extracted using the Qiagen Dneasy Plant Pro kit. A modification to the lysis step was performed by snap-freezing the sample, having been transferred to the disruption tubes provided in the kit, with liquid nitrogen. Afterwards 500µL of CD1 buffer (provided by the kit) was added and the tubes were disrupted using a FastPrep-24™ 5G Bead Beating Grinder and Lysis System (5.5 m/s x 30 sec). 20µL Proteinase K (20mg/ml) was then added to each sample and then disrupted again using the Fastprep gear, this time 5.5 m/s x 15 sec. Samples were incubated at 56°C overnight and then processed following the Qiagen Dneasy Plant Pro kit protocol. Whole-genome shotgun sequencing was performed on the DNA samples using an Illumina NovaSeq6000 targeting 10 Gb of 2×150 PE data per sample.

### Population genetic structure, F_ST_-Outlier Detection, phylogenetic tree

Reads were filtered and trimmed using the paired-end function of Trimmomatic v0.39 [28], before being mapped to the *Asterionella formosa* reference genome (NCBI accession: GCA_002256025.1) using the package bwa v0.7.17. We then removed optical and PCR duplicates using the package Picard v2.7.1 and called variants and invariants sites using the function samtools mpileup v1.10. We performed a subsequent filtration of the data removing duplicates, indels or entries which quality score did not match our cutoff values (%QUAL<20, INFO/DP>15). Reads passing the filtering steps were used for downstream analyses.

To identify subpopulations within the *A. formosa* isolates we used the Bayesian clustering analysis software STRUCTURE v2.3.4 [29,30]. We set the number of genetic clusters (*K*) to range between 1 and 10 and performed three independent runs for each *K* using a burn-in of 50,000 iterations and a run length of 250,000 iterations. *K* values between 7 and 10 did not converge, despite futher extension of the chains to 450,000 iterations, hence we evaluated *K* values between 1 and 6. To infer support for the optimal number of subpopulations we calculated *ΔK* using the Evanno method implemented in the program STRUCTURE HARVESTER v0.7 [31].

We detected *F_ST_*-outliers using the fsthet v1.0.1 package in R, which identifies outliers among genome-wide SNPs generating smoothed quantiles for the *F_ST_*-heterozygosity distribution [32]. All analyses in R were performed using R and Rstudio v4.3.2 [52]. To infer which genes contained outlier loci and the putative functions of their proteins, we performed iterative gene model prediction of the *A. formosa* reference genome using MAKER v3.01.04 [33]. Functional annotations were made using the predicted protein set from the fourth round of gene model inference using eggNOG-mapper v2.1.12 [34].

We constructed a phylogenetic tree with IQ-TREE v2.2.2.6 [35], applying the generalized time-reversible (GTR) model of nucleotide substitution [36] correcting for ascertainment bias inherent to SNP data (+ASC) [37]. Topology support was inferred using the ultrafast bootstrap method with 1000 replicates [38]. The consensus tree was then visualized in R using the package ggtree v3.10.0 [39]. Nucleotide diversity (π) between genotypes was calculated according to [40] using the software DnaSP v6. Synonymous and nonsynonymous mutations were identified by comparing the variant coordinates in the VCF files with the exons in the GFF file produced from MAKER, and evaluating the impact that the variant of interest has on AA substitution using the software SnapGene v7.1.1.

### Traits measurements and analysis

We conducted experiments to assess functional traits using seven *Asterionella* genotypes picked randomly from March (31.03.21) and June (30.06.21). For each genotype we estimated maximum growth rates (µ_max_), carrying capacity (*K_c_*), half-saturation constant of P (P-K) and Si (Si-K), sinking rate and fitness (AUC, area under the curve [41]) in different environments. All traits were derived performing growth experiments. Culture preparation and experimental conditions varied depending on trait, but in general, all experiments were conducted in culturing flasks with a final volume of 45 mL. Cultures were placed on orbital shakers at 80 rpm and the light intensity was kept constant at 45 µmol m^−2^ s^−2^ (PAR). Cultures were grown with three replicates per isolate, and 3×200 µL of each replicate were sampled directly to a microwell plate every 24/48 hours (see below) until stationary phase was reached. The microwell plates were immediately fixed with Lugol solution (5% final concentration) and stored at 4°C in the dark until measurement. Growth was monitored using a high content microscope (ImageXpress® Micro 4 High-Content Imaging System, IXM). We acquired four images for each well, covering the entire well surface, using a Cy5 filter set (algal autofluorescence, 642 nm) under a 5x magnification. For the initial and last days, we additionally acquired images with transmitted light to analyze cell morphology. The images were analyzed using the software MetaXpress® High Content Image Acquisition and Analysis and growth curves estimated from the cells total fluorescent area (hereafter density) as a proxy for cell count. The mean densities of the three subsamples taken per replicate were used to calculate the growth rates of each isolate replicate. We estimated growth rates comparing a linear and a logistic growth model [42,43] via AIC (table S1), calculated by fitting the models to each combination of isolate and treatment on log-transformed densities. Afterwards, using the predicted growth curves from the best-fitting model, we computed linear regressions for each isolate and treatment over four consecutive time steps (i.e., 1-4 days, 2-6 days depending on the experiment, see below) to find the per-capita µ_max_.

A principal component analysis (PCA) was constructed from the trait data to assess the differences in phenotypes among *Asterionella* isolates. The input traits were standardized prior to PCA analysis to account for differences in units of measurement. Because of measurement errors in the Si experiment, we had to remove 2 replicates (*Af_8* rep3, *Af_15* rep 1) from the PCA analysis. To extract the contributions of each trait to the PC-axes we used the R package factoextra v1.0.7[44] and to test for trait covariation we calculated pairwise Spearman’s rank correlations, which we then plotted in a trait correlation matrix using the corrplot v0.92 package in R [45]. Finally, to test if season had a significant effect on isolate distribution in the PCA, we performed a PERMANOVA using the function adonis2 from the vegan package v2.6-4 [46].

### Maximum growth rates and carrying capacity

To assess µ_max_ and *K_c_* of *A. formosa*, each isolate was grown in triplicates for eight days at 20°C under constant light. Samples were taken daily and flasks re-shuffled on the shaker after each sampling to randomize the flask layout. Because of measurement errors three out of the 72 subsamples of day1 were discarded from the dataset. We estimated µ_max_ for each isolate over the first four days and found the logistic model to be the best fit for all isolates using AIC (table S1a). We obtained values for *K_c_*directly from the logistic growth model.

### Temperature optimum

To estimate the temperature optima (T_opt_), we tested the isolates at seven different temperatures: 5, 10, 15, 20, 22.5, 25, 30°C, which were chosen according to previous experiments on *Asterionella formosa* [47,17] and, except for 30°C, encompass the yearly natural water temperatures in Lake Constance. Before the experiment all the cultures were acclimated to the different temperatures for one – three weeks to account for differences in generations times at different temperatures [48]. The experiment lasted from a minimum of 8 days (temperatures 25 and 30°C) to a maximum of 18 days (5°C) depending on culture growth and mortality. During the experiment samples were taken daily and cultures re-shuffled on the shaker after each sampling. Isolates did not survive at 25 and 30°C, therefore we considered a growth rate equal to 0 when quantifying the thermal performance curves (TPCs). We removed day7 at 22.5°C from the analysis because the plate went lost. The mean densities of the isolates were used to calculate growth rates over the first four days, from which we then computed TPCs and obtained the T_opt_ using the R packages *rTPC* and *nls.multstart* [49]. We compared the fit of 16 different TPC via AIC comparison, yielding *Sharpeschoolfull 1981* [50] as the best-fitting for all isolates (table S1c).

### Nutrient half-saturation constant

To estimate half-saturation constants of phosphorus and silicate, isolates growing at exponential phase were transferred to 15 mL centrifuge tubes and centrifuged at 1972 x g for 5 min at 20°C. Cell pellets were washed twice with 10 mL of WC medium without PO_4_ or Si-SiO_2_ (WC 0 P and WC 0 Si), accordingly. The pellet was resuspended in 2 mL WC 0 and transferred to flasks with WC 0 P or WC 0 Si in a final volume of 40 mL. Cultures were then grown for 10 days at 20°C in WC 0 P to assure P depletion. Since *A. formosa* does not store silica, the cells were starved in WC 0 Si for 4 days [51].

We grew the isolates at 1, 2, 4, 8, 12, 20 µM Si-SiO_2_ and at 0.032, 0.081, 0.32, 0.64, 1.61, 50 µM PO_4_, concentrations that span limiting to saturating levels [52]. All other elements in the medium were not manipulated. In all cases, samples were taken every second day and flasks were randomly shuffled on the shaker post-sampling. Because of measurement errors mainly due to low initial densities, we had to remove 12 out of 3150 values from the Si experiment and four out of 3273 from the P experiment before estimating half-saturation constants. We estimated µ_max_ for each isolate-nutrient combination over the days 2-6 using the best-fitting model chosen by comparing AICs (table S1b) and we then fit the Monod equation to obtain half-saturation constants [53]:

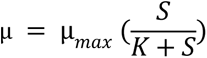

### Sinking rate

To assess the sinking rate of *A. formosa*, the isolates were sampled three times from one replicate during the exponential phase. 5 µL of each culture were pipetted into a well of a 48-well plate containing 995µL of medium to reach a final volume of 1 mL. Before conducting the sinking assay, the plate was mixed at 650 rpm x 7 minutes using a plate vortex to resuspend all cells. Images were then acquired at the bottom of the plate every 10 minutes for 11 time points using the high content microscope IXM. We set-up this technique according to the standard SETCOL method with the advantage of avoiding handling large volumes of cultures [54].

Sinking rate was calculated as follow [54]:

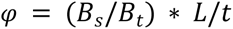

where φ is sinking rate, *B_s_* is the biomass settled at the bottom of the well, *B_t_* is the total biomass in the well, *L* is the sample depth and *t* is the settling interval.

### Fitness in different environments

We estimated fitness as Area Under the Curve (AUC) in different environments by manipulating simultaneously temperature and nutrient content of the culture media to simulate the conditions of Lake Constance in March and June. The temperatures used were 5 and 20°C. The nutrients concentrations used at 5°C were 20 µM Si-SiO_2_ and 0.162 µM PO_4_; while at 20°C were 4 µM Si-SiO_2_ and 0.162 µM PO_4_. Cultures were cleaned and starved before conducting the experiment by growing the samples for eight days in WC 0 P and, subsequently, four days in WC 0 Si, although adding a small amount of PO_4_ (0.064 µM) to prevent cell death. We took samples every second day and we estimated fitness as AUC for each isolate-treatment combination using the R package DescTools v0.99.53 [55]. AUC values were divided by cell length to account for differences in isolate sizes.

## Results

### Spring ecological dynamics

The spring phytoplankton dynamics in 2021 were consistent with the seasonality described by the PEG model (figure 1). Algal biomass increased from March to May along with thermal stratification and decreasing nutrient availability (figure 1B,D) with diatoms (Pennales + Centrales) contributing the most to the total biovolume with an average relative biovolume of 52.24% ± 24.33% (figure 1C). Small Centrales covered the highest proportion of diatom biovolume in early spring (average 39.33 ± 6.92 %), while large Pennales were dominant in late spring (average 48.53 ± 14.47 %). Among Pennales, *Asterionella formosa* covered the middle ranks of the phytoplankton rank-biovolume distributions from March to May, stepping up to the top ten rank positions in June (figure S1).

**Figure 1.**
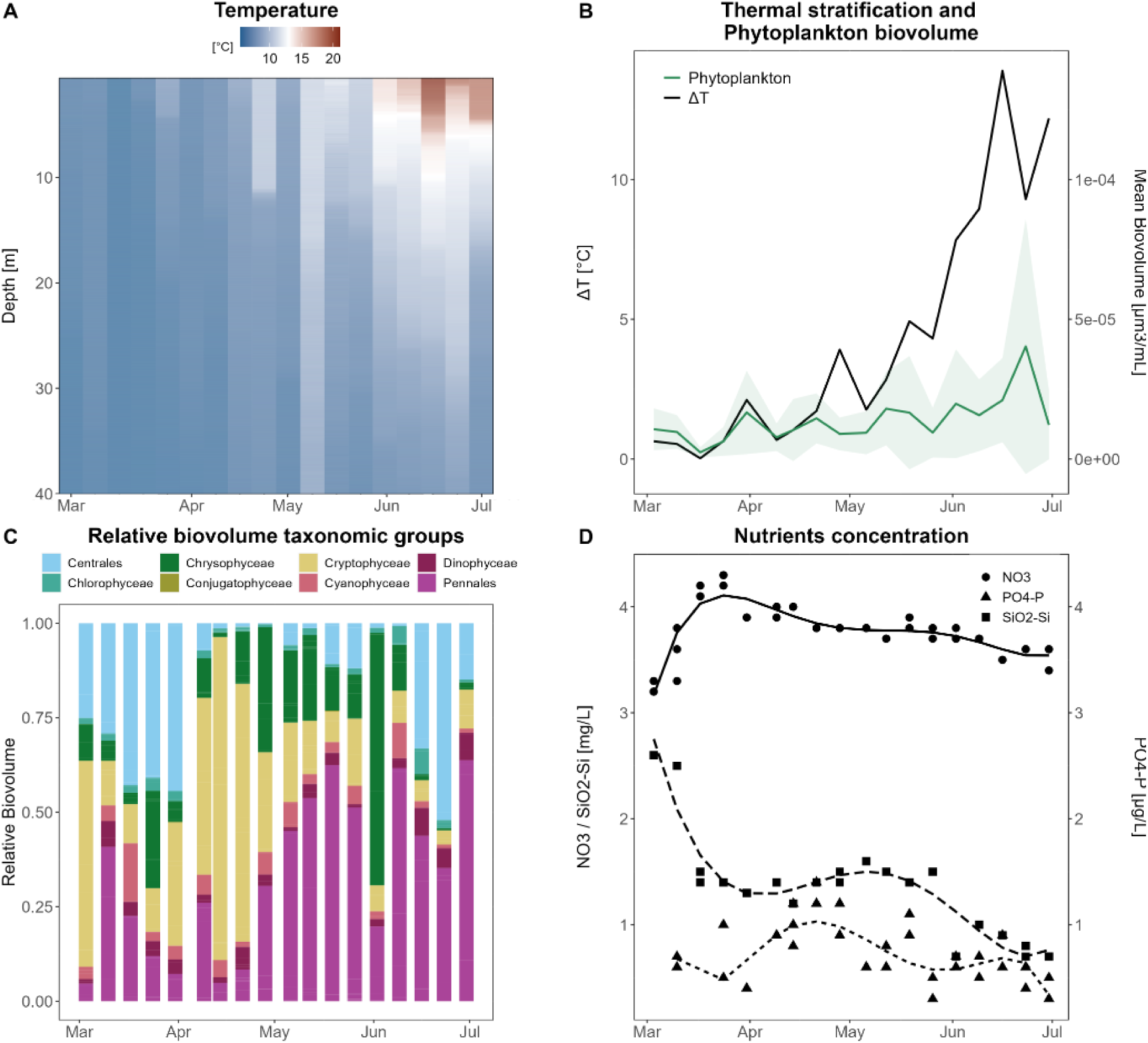
Temporal changes in environmental parameters and phytoplankton community in Lake Constance along the spring bloom 2021. A) Temperature profiles. B) Thermal stratification (ΔT) and phytoplankton development. The green shaded area represents the 95% confidence intervals. C) Contribution of the different taxonomic groups to the total phytoplankton biovolume. D) Nutrient concentrations. Point and line shapes correspond to the different nutrients measured. Lines are fitted polynomial regressions.

### Asterionella genotypic diversity

2917 polymorphic sites were present in the 33 isolates after filtering for sites present in at least 90% of the isolates. Each sample was a unique genotype in the SNP matrix; hence, we did not sample identical clones representing separate isolates. The clades recovered on the phylogenetic tree displayed a strong seasonal correlation (figure 2A). The total nucleotide polymorphism (π_T_) calculated between all isolates was 0.079 ± 0.001 (S.D.), while the nucleotide diversity estimated according to seasonality was equal to 0.092 ± 0.002 (S.D.) in early spring and to 0.100 ± 0.003 (S.D.) in late spring (figure 2B). These results suggested that genetic polymorphism was higher in late spring compared to early spring.

**Figure 2.**
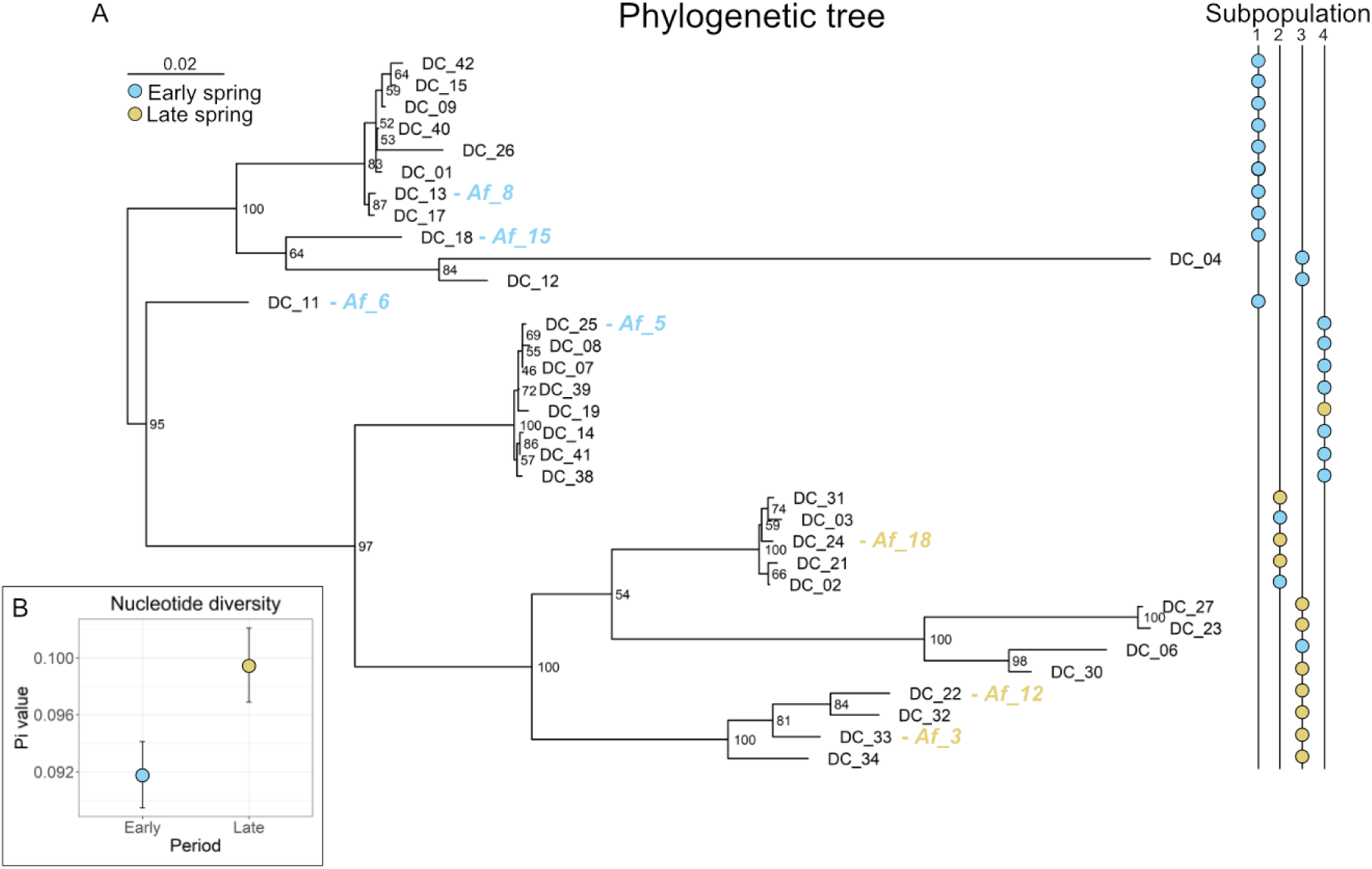
A) Phylogenetic tree of *Asterionella* isolates. Colors correspond to the seasonal phase: blue = early spring, yellow = late spring. Node support values obtained using ultrafast bootstrap are reported as node labels. Blue and yellow text labels following the tip labels correspond to samples used in the phenotypic assays. B) Nucleotide diversity between early and late spring.

The analysis of genetic structure partitioned the isolates into 4 different genetic clusters or subpopulations (figure 3). The subpopulations’ contributions to the overall population for a given period showed a temporal shift, with subpopulation one dominating in early spring (45.45% of the total proportion) and subpopulation three dominating in late spring (63.64% of the total proportion). Subpopulation two was predominant in late spring (27.27%), while subpopulation four in early spring (31.82%).

**Figure 3.**
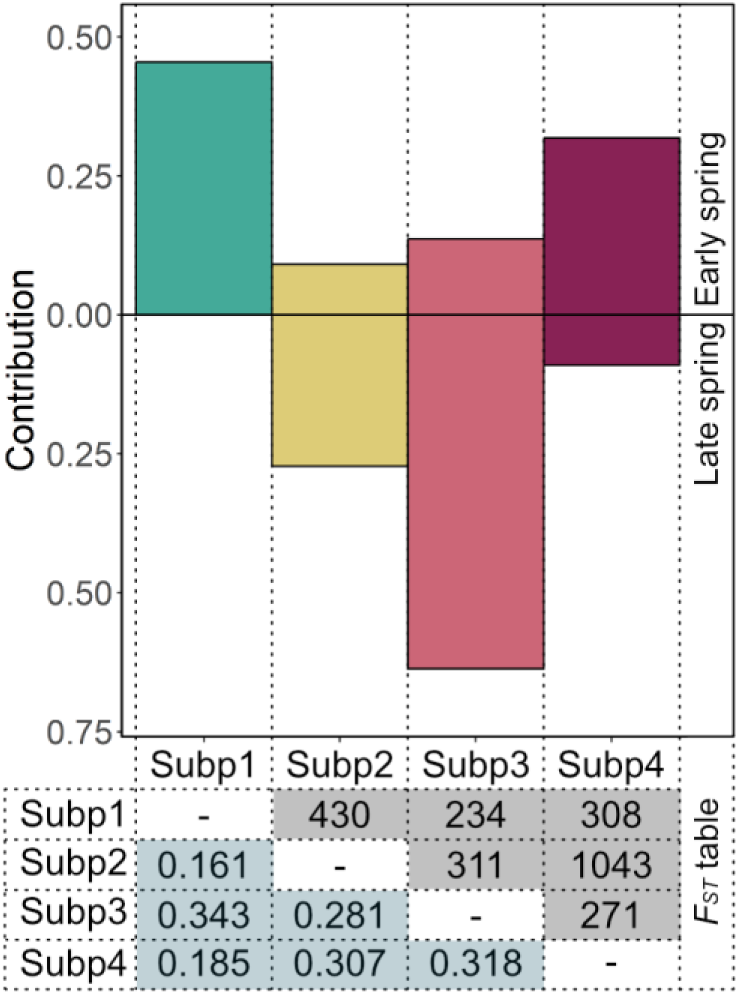
Contribution of the different *Asterionella* subpopulations to early and late spring. The table below reports in grey the average *F_ST_* values and in blue the number of *F_ST_* outliers between the different pairs of subpopulations (Subp).

We then estimated pairwise *Fst* values for each possible subpopulation combination. The average *F_ST_* values ranged from a minimum of 0.161 to a maximum of 0.343 (figure 3). The highest population differentiation was found between subpopulations one – three (*F_ST_:* 0.343) and three – four (*F_ST_:* 0.318); the lowest differentiation in the pairs one – two (*F_ST_:* 0.161) and one - four (*F_ST_:* 0.185). A total of 1576 outlier loci were detected among all pairwise combinations using fsthet (figure S4). Only two of the 1576 outliers were found in all combinations, three in five pairwise combinations and 37 in four pairs. We found in total 233 *F_ST_* outliers between subpopulations one and three, of which 68 were identified as being under balancing selection and 165 under positive selection by fsthet. Nine of these 233 sites corresponded to loci within exons for which the parent genes had functional annotations (table S2). Four of the loci mapped to genes involved in defense/immune responses (BAM3, MIK2, NLRC3, NLRC3) and, except one (NLRC3), all of them showed positive selection. The same pattern was found between pair two – four, with two additional sites (NPR4, At5g63930) under positive selection involved in basal defense against pathogens. The two clusters which contributed the most to the early period, cluster one and four, showed instead an overall balancing selection in loci involved in immune response against pathogens (table S2). The substitutions in amino acid sequences of the loci involved in immune response were, excepts for few cases for NPR4 and BAM3, nonsynonymous (table S2). We found this pattern to correlate to the seasonal parasite epidemic events observed in the lake samples: we detected no chytrid attached to *Asterionella formosa* in early spring, when algal densities were low (average 2.2 cells/mL in March-April; figure 4). On the contrary, in June we observed relatively high numbers of chytrids per *Asterionella* colony, with a maximum of 6.38 chytrids/colony when the diatom also had its highest densities (average 60.4 cells/mL in May-June).

**Figure 4.**
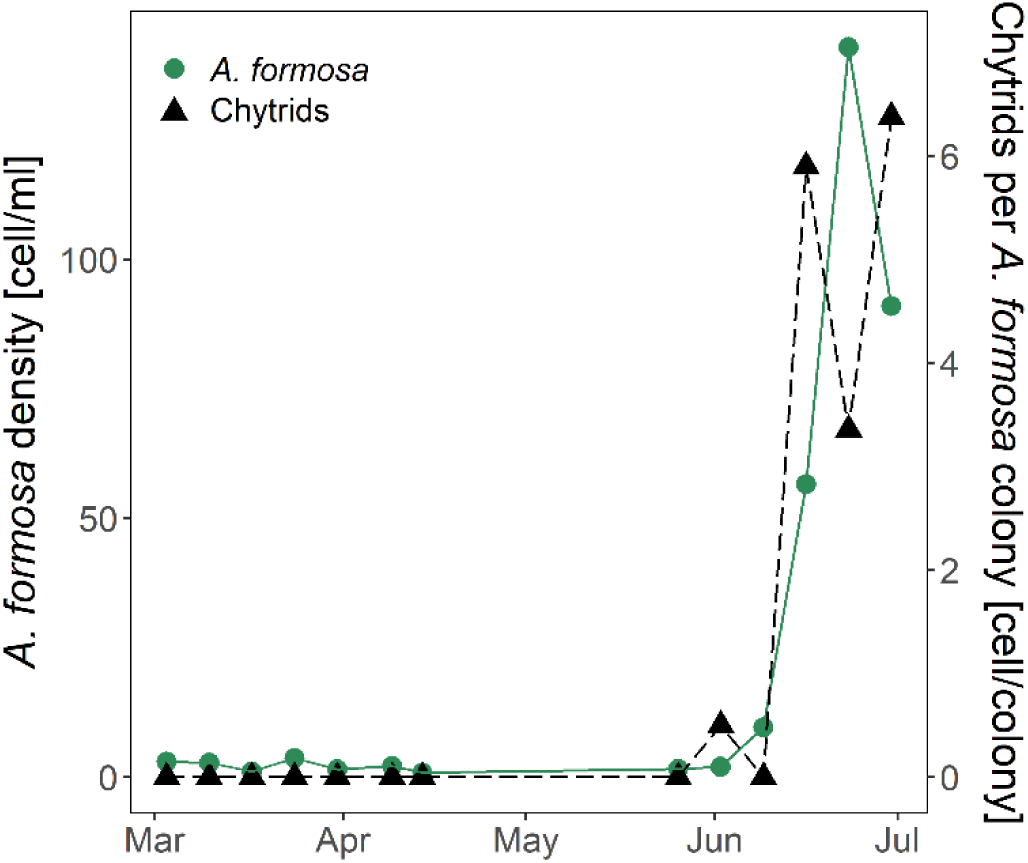
Densities of *A. formosa* and chytrids in spring 2021.

### Asterionella *phenotypic diversity*

Among the 33 unique genotypes identified, we chose randomly a subset of seven to assess if genetic diversity also has consequences on traits and fitness. Four of the genotypes were isolated in March (31.03.21) and the other three in June (30.06.21), representing early and late spring. PCA analysis uncovered a seasonal phenotypic variability among the *Asterionella* genotypes (figure 5). We did not include T_opt_ in the PCA since information for two genotypes was missing. The average T_opt_ found across the genotypes tested was 21.69 ± 0.20 (S.D.) °C (table S3). The two main axes of the PCA explained 60.5% of the total variation. The different genotypes clustered between periods along PC1, which accounted for 34.5% of the total variation and was mostly explained by the traits: *K_c_* (24.00%), sinking rate (18.46%), Si-K (18.40%), P-K (11.48%) and µ_max_ (11.07%), and by fitness at 20°C (AUC_20: 14.21%). Length (30.25%) and AUC_5 (28.73%) and AUC_20 (14.08%) characterized PC2. Among certain traits and trait - season combinations we detected consistent correlations (figure S3). Specifically, µ_max_, *K_c_* and length correlated positively with June genotypes (*ρ* =0.45, *P* = 0.05; *ρ* =0.87, *P =* 1.65e-06 and *ρ* =0.52, *P =* 0.02). µ_max_ and *K_c_* had a negative relationship with P-K (*ρ* = −0.58, *P* = 9.49e-03 and *ρ* = −0.54, *P =* 0.02), *K_c_*correlated significantly with µ_max_ (*ρ* = 0.49, *P =* 0.03) and Si-K with sinking rate (*ρ* = 0.58, *P =* 0.01). Regarding fitness, AUC_20 exhibited a negative significant correlation with Si-K (*ρ* = −0.57, *P =* 0.01) and AUC_5 showed a positive relationship with length (*ρ* = 0.68, *P =* 0.002). Fitness in the two different environments did not correlate with seasonal phase (*P >* 0.05). Taking in account all phenotypic traits measured and fitness, the isolates clustered significantly according to seasonal phase in the PCA (PERMANOVA test, 1000 permutations, *P* = 9.99e-04)

**Figure 5.**
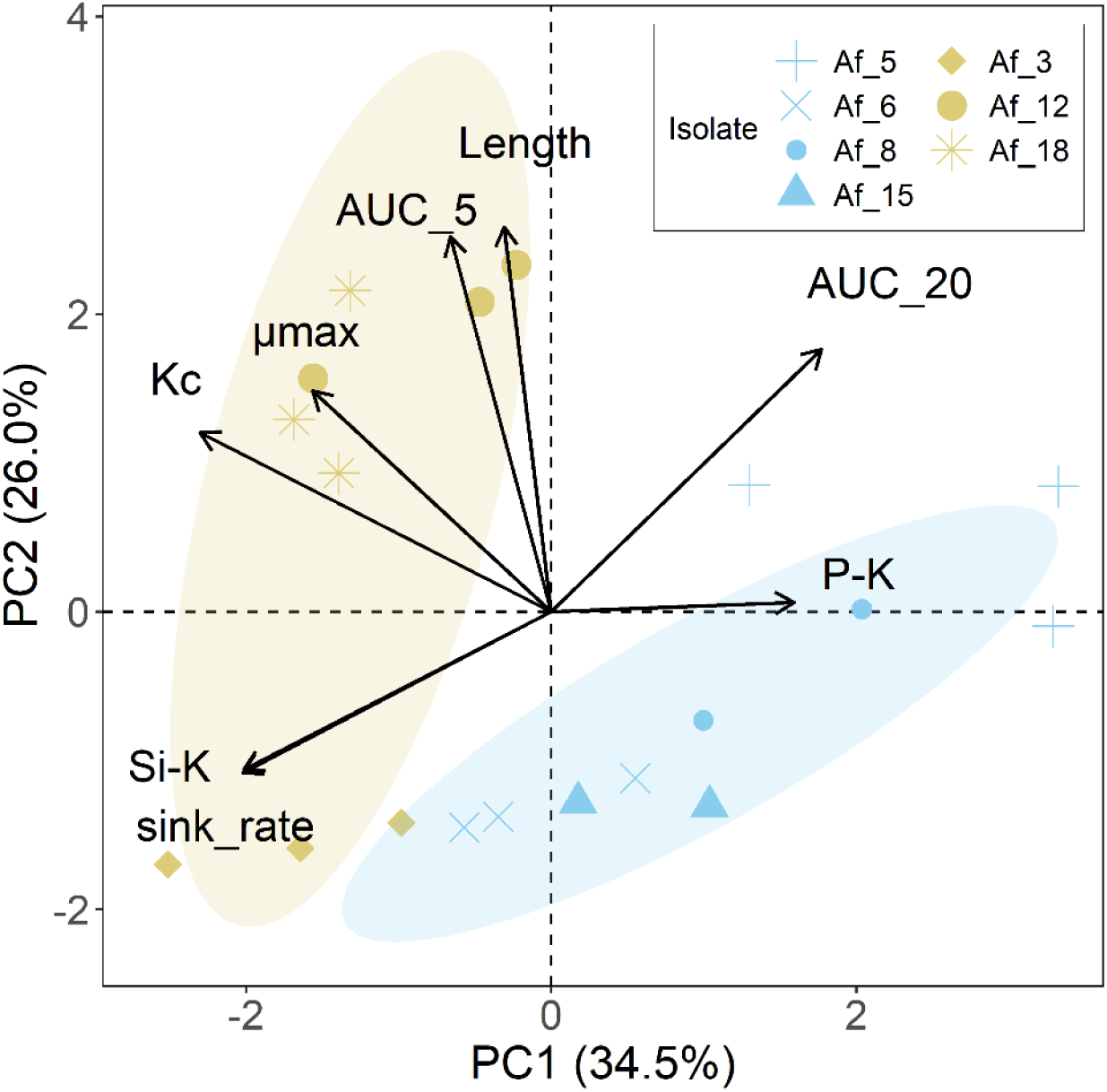
PCA analysis performed with 8 different measured traits for 7 *Asterionella formosa* genotypes. The colors represent the period (blue = early spring, yellow = late spring) and shapes the different genotypes.

## Discussion

Within this study we aimed to investigate eco-evolutionary dynamics in a natural system in the presence of seasonal changes. Our results showed that the *Asterionella* population underwent seasonal genotypic and phenotypic shifts through the spring season of 2021. The genetic structure of the *Asterionella* population revealed a high genetic diversity within the lake (figure 2), with the highest differentiation found between subpopulation pairs one – three (*F_ST_*: 0.343) and three – four (*F_ST_*: 0.318) (figure 3B). This can be explained by the fact that subpopulations one and four appeared mostly in the early period while subpopulation three had the highest contribution in late spring, suggesting a shift in genotype frequency over time, similar to the seasonal shift reported for *Daphnia mendotae* clones in Lake Oneida [14]. Among sites under selection between pairs one – three, three – four and additionally two – four we found that genes involved in defense/immune responses against pathogens are generally under positive selection (table S2). This suggested that there is most likely selection on genes involved in pathogen resistance between *Asterionella* genotypes of early and late season. This pattern correlated to the chytrid infection rates. We did not detect any parasite infection in early spring, when *Asterionella* was present in low numbers. Instead, during the *Asterionella* bloom in June, colonies were highly infected. Our observation is consistent with previous findings which showed that parasitic chytrids follow the seasonal dynamics of the environment and their hosts [57,58] and suggested that parasite population growth is often host-density dependent [18,56,59]. Additionally, we found nucleotide polymorphism to be higher in late compared to early spring, supporting the expectation that within blooming phytoplankton populations intraspecific variation is high [60–62] and that high-parasite pressure leads to an increase in host genetic diversity, since parasitism is often genotype-specific [18,57] and higher diversity favours higher resistance [56,63].

The genetic structure of *Asterionella* population could predict the seasonal clusters detected in the phenotypic multitrait space, with genotypes from the same seasonal phase being phenotypically more similar (figure 5). While the genetic analyses implicate the importance of parasite population dynamics throughout the season, seasonality in traits was mostly defined by PC1 (*K_c_*, Si-K, sinking rate, AUC_20, P-K and µ_max_) in the PCA. Length distribution varied along spring (figure S2), with a March cell size that was on average smaller than in June [64]. Length was not correlated to either P-K or Si-K (*P* > 0.05) in our study, however factors other than size might be influencing cell physiology and nutrient affinity [65,66]. Few studies reported that large-cell *Asterionella* showed higher growth rate than small-cell colonies [64], consistent with our findings for fitness.

As required for eco-evolutionary dynamics we here showed seasonal changes in traits and genotypes, however, even though the isolates were partitioned according to seasonal phase and we detected some trait-fitness correlations, we did not find fitness variations to be linked to season. The trait-fitness correlations we observed are the relationship between AUC_20 and Si-K, which means that a higher competitive ability for Si correlates with higher fitness at 20°C in Si-limiting environment; and the correlation between AUC_5 and length, which translates into higher fitness at 5°C and low P concentrations when cells are larger. One explanation why we did not find correlations between fitness and season could be the similar T_opt_ (21.69 ± 0.20 S.D.) and temperature range observed for all strains. Furthermore, P-K did not correlate with season and PO_4_-P concentrations were low along the entire spring in the lake (figure 1). Si-K was for all genotypes lower than lake Si concentration (minimum concentration: 11.83 µmol/L) and *Asterionella* did likely not experience Si limitation. This indicates that temperature and nutrients are probably not a major driver of *Asterionella* seasonal differentiation and consequently explains why the genotype fitness tested in different abiotic conditions is not linked to season. Biotic stressors might be the selective force underlying the evolutionary changes observed.

The overall high genetic diversity of the *Asterionella* population is consistent with former studies on phytoplankton which showed that populations can be highly genetically differentiated [16,67,60]. This might indicate that asexual proliferation is predominant and maintained for long time spans in phytoplankton populations [16,21]. The *F_ST_* values found in our study, as well as the ones reported in the literature for *Asterionella* [16], are in a range in which researchers typically define cryptic species for diverse organisms, like ascidians (*F_ST_*: 0.276) [68], or even within some diatoms species (*F_ST_*for *Pseudo-nitzschia pungens*: 0.481) [69]. Population genetics studies on diatoms need to further investigate, whether we are facing large intraspecific genetic differentiation or a potential ecological speciation in the system [16,70].

The high level of genetic and phenotypic differentiation among *Asterionella* isolates found in this work offer an insight on intraspecific dynamics of natural phytoplankton populations in response to a changing environment over time and add to trait-based ecology that population phenotypic and genetic diversity can be as important as interspecific variation in explaining community dynamics [71,72]. We observed how ecological dynamics, in our case the presence of a parasite, likely contributed to generate genetic shifts in a population to allow for local adaptation on a seasonal time scale. This example add to recent work [14] of how eco-evolutionary dynamics are critical for natural plankton populations to rapidly adapt to seasonal fluctuations. We therefore highlight the importance of including eco-evolutionary processes in the already existing models when studying seasonality, to be able to better understand population and ultimately community dynamics in natural systems.

## Data accessibility

The data presented in this study can be found in the Dryad Digital Repository: doi:10.5061/dryad.k6djh9wfc.

## Authors’ contributions

D.C and L.B: conceptualization, writing—review and editing; D.C: sampling, formal analysis, molecular analysis, methodology, data curation; J.B.K: molecular analysis, methodology, supervision, writing—review and editing; P.M: phytoplankton and chytrids counting; L.B: project funding, supervision.

## Acknowledgements

We thank A. Sulger, D. Schleheck and J. Schmidt for technical assistance in the field and the student assistants, Maximiliane Ziefle and Riya Mathur, who helped in the nutrient meaurements and culture maintainance in the laboratory.

## Conflict of interest declaration

The authors declare no competing interests.

## Funding

This research was done as part of the Research Training Group (RTG)—R3 (Responses to biotic and abiotic changes, Resilience and Reversibility of lake ecosystems) funded by the Deutsche Forschungsgemeinschaft (DFG, German Research Foundation— 298726046/GRK8872). This work was funded by the Gordon and Betty Moore Foundation to LB (grant #9196; https://doi.org/10.37807/GBMF9196). This material is based upon work supported by the NSF Postdoctoral Research Fellowships in Biology Program under Grant No. 2109477, awarded to JBK.

## Notes

### Competing Interest Statement

The authors have declared no competing interest.

