## Supplementary material for "Seasonal genotypic and phenotypic differentiation of a cosmopolitan freshwater diatom"

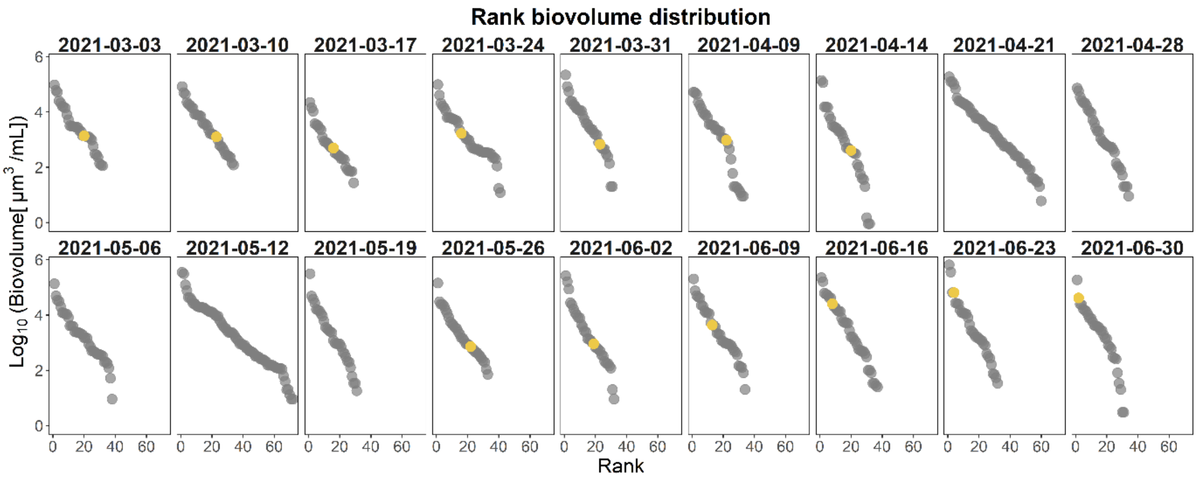


**Figure S1.** Rank biovolume distribution plots. Each dot represents one species, yellow dots represents *Asterionella formosa*.

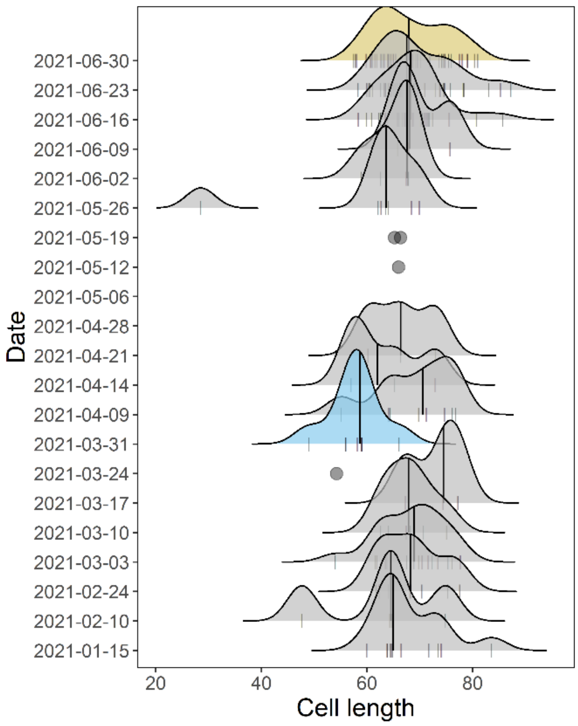


**Figure S2.** Cell length distribution of *Asterionella formosa* during the spring bloom 2021. Yellow and blue represent the dates in March and June in which the samples for the phenotypic assays were isolated.

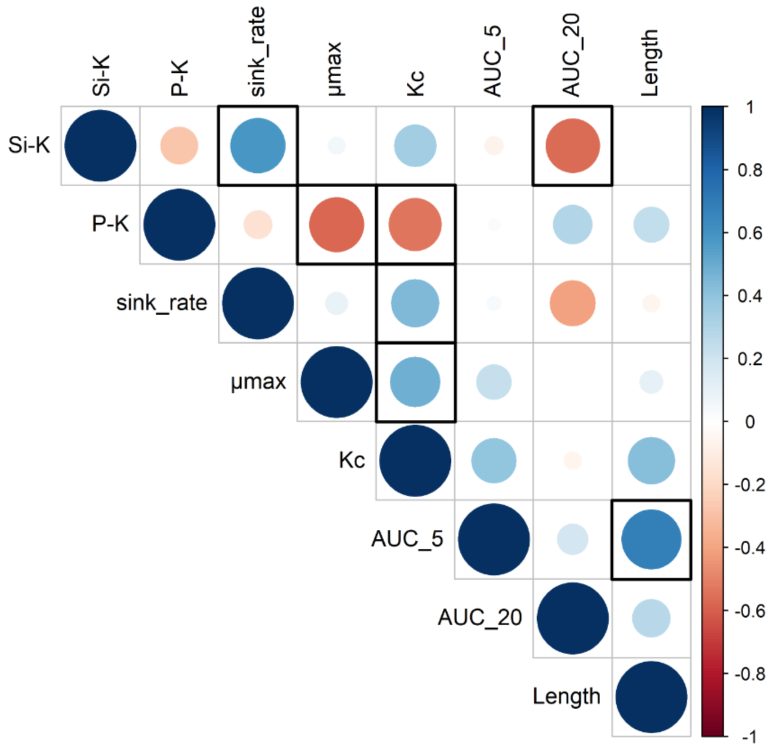


**Figure S3.** Correlation plot of *Asterionella* trait data. Correlations are estimated based on Spearman's rho. We highlighted borders of significant correlations ($P < 0.05$).

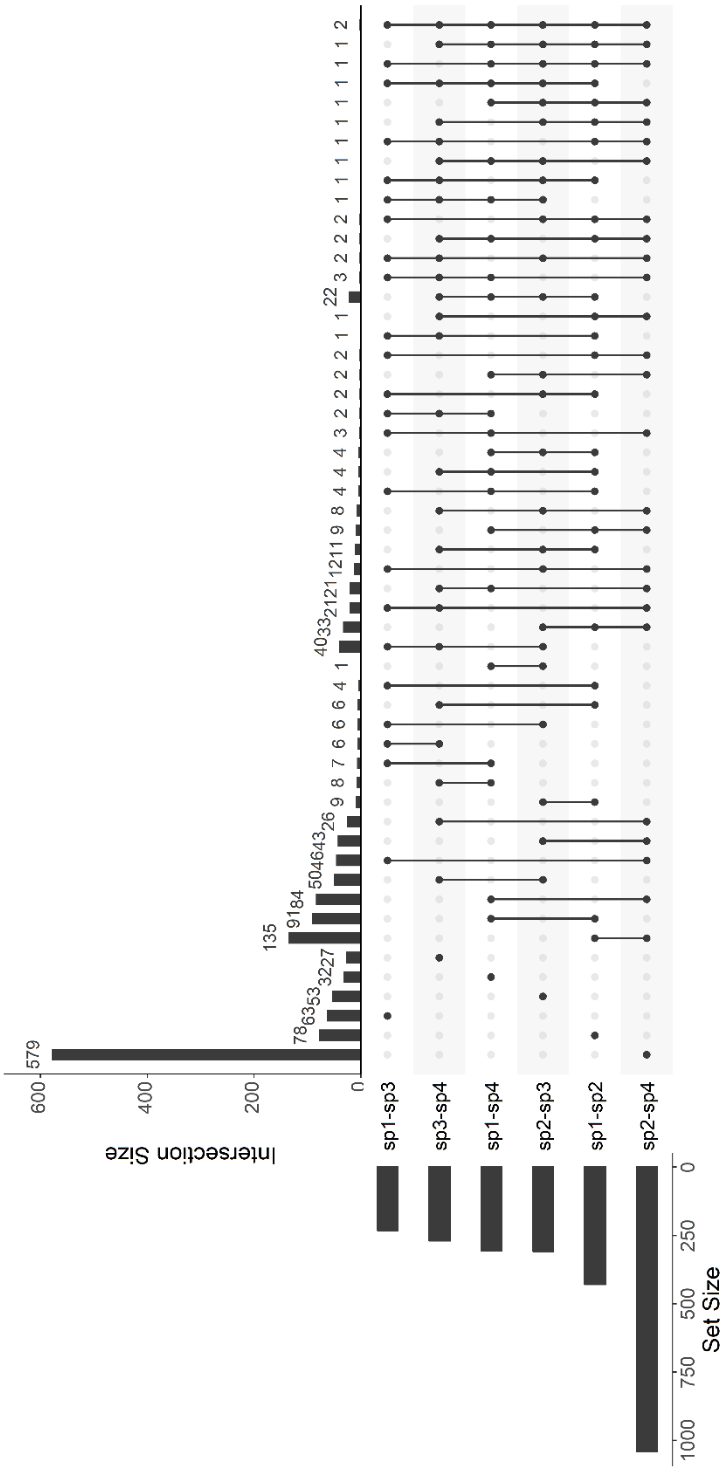


**Figure S4.** Intersection plot. Visualization of the number of $F_{ST}$ outliers found and shared between pairs of subpopulations. The set size defines the number of outliers per pair of subpopulations, the intersection size describes how many outliers are present per pair intersection. The plot was obtained using the package UpSetR [1].

**Table S1.** AIC comparison of the fitted models for the different experiments. a) Growth experiment; b) P and Si half-saturation constant experiments; c) Temperature experiment. For the TPCs, we fitted the following models (1-16): Boatman_2017, Briere2_1999, Gaussian_1947, Joehnk_2008, Lactin2_1995, Oneill_1972, Ratkowsky_1983, Thomas_2012, Weibull_1995, Sharpeschoolhigh_1981, Sharpeschoolfull_1981, Delong_2017, Beta_2012, Deutsch_2008, Flinn_1987, Pawar_2018 [2].

**a)**

| Growth experiment | | | |
|---|---|---|---|
| **Isolate** | **Rep** | **AIC linear model** | **AIC logistic model** |
| Af_5 | Rep1 | 19.22 | **5.43** |
| | Rep2 | 18.32 | **1.08** |
| | Rep3 | 20.10 | **3.07** |
| Af_6 | Rep1 | 22.47 | **6.60** |
| | Rep2 | 22.20 | **7.70** |
| | Rep3 | 21.92 | **5.92** |
| Af_8 | Rep1 | 20.47 | **5.73** |
| | Rep2 | 22.15 | **4.49** |
| | Rep3 | 21.37 | **7.56** |
| Af_15 | Rep1 | 20.30 | **4.84** |
| | Rep2 | 20.08 | **5.15** |
| | Rep3 | 20.10 | **8.68** |
| Af_3 | Rep1 | 19.84 | **9.91** |
| | Rep2 | 19.65 | **8.08** |
| | Rep3 | 20.32 | **7.96** |
| Af_12 | Rep1 | 22.61 | **8.38** |
| | Rep2 | 22.55 | **7.06** |
| | Rep3 | 21.74 | **6.81** |
| Af_18 | Rep1 | 25.01 | **5.88** |
| | Rep2 | 25.23 | **10.80** |
| | Rep3 | 24.52 | **7.73** |

**b)**

| **P half-saturation experiment** | | | | | **Si half-saturation experiment** | | | | |
|---|---|---|---|---|---|---|---|---|---|
| **Isolate** | **Rep** | **Treatment [µM $PO_4$]** | **AIC linear model** | **AIC logistic model** | **Isolate** | **Rep** | **Treatment [µM Si-$SiO_2$]** | **AIC linear model** | **AIC logistic model** |
| Af_5 | Rep1 | 0.032 | -10.86 | **-19.46** | Af_5 | Rep1 | 1 | **-2.13** | NA |
| | Rep2 | | -0.77 | **-6.61** | | Rep2 | | 3.11 | **-2.46** |
| | Rep3 | | -9.36 | **-12.31** | | Rep3 | | 1.87 | **0.70** |
| Af_5 | Rep1 | 0.081 | -8.06 | **-9.03** | Af_5 | Rep1 | 2 | 9.95 | **3.66** |
| | Rep2 | | -5.20 | **-8.15** | | Rep2 | | 8.05 | **-3.13** |
| | Rep3 | | -2.47 | **-11.06** | | Rep3 | | 6.23 | **-8.00** |
| Af_5 | Rep1 | 0.32 | 18.85 | **-6.42** | Af_5 | Rep1 | 4 | 16.04 | **-2.65** |
| | Rep2 | | 19.30 | **-2.77** | | Rep2 | | 19.30 | **-3.85** |
| | Rep3 | | 22.28 | **-2.01** | | Rep3 | | 20.15 | **2.23** |
| Af_5 | Rep1 | 0.64 | 24.72 | **-5.78** | Af_5 | Rep1 | 8 | 22.69 | **2.80** |
| | Rep2 | | 24.70 | **5.36** | | Rep2 | | 22.58 | **0.45** |
| | Rep3 | | 24.15 | **-3.59** | | Rep3 | | 24.92 | **2.84** |
| Af_5 | Rep1 | 1.61 | 21.10 | **12.00** | Af_5 | Rep1 | 12 | 22.32 | **1.83** |
| | Rep2 | | 20.75 | **14.98** | | Rep2 | | 24.17 | **4.82** |
| | Rep3 | | 17.86 | **8.14** | | Rep3 | | 23.13 | **5.00** |
| Af_5 | Rep1 | 50 | 21.46 | **13.11** | Af_5 | Rep1 | 20 | 24.91 | **4.76** |
| | Rep2 | | 21.66 | **17.57** | | Rep2 | | 24.54 | **2.94** |
| | Rep3 | | 21.37 | **17.59** | | Rep3 | | 24.24 | **2.61** |
| Af_6 | Rep1 | 0.032 | -0.05 | **-12.01** | Af_6 | Rep1 | 1 | 6.79 | **1.14** |
| | Rep2 | | -0.83 | **-6.41** | | Rep2 | | 0.36 | **-0.01** |
| | Rep3 | | 4.12 | **-1.08** | | Rep3 | | 6.05 | **4.03** |
| Af_6 | Rep1 | 0.081 | 3.60 | **-11.01** | Af_6 | Rep1 | 2 | 5.34 | **-3.63** |
| | Rep2 | | 5.96 | **-7.33** | | Rep2 | | 5.51 | **3.47** |
| | Rep3 | | 5.74 | **-9.60** | | Rep3 | | 9.92 | **7.61** |
| Af_6 | Rep1 | 0.32 | 20.69 | **-1.85** | Af_6 | Rep1 | 4 | 11.80 | **-0.45** |
| | Rep2 | | 20.53 | **2.83** | | Rep2 | | 14.24 | **-18.84** |
| | Rep3 | | 22.77 | **4.81** | | Rep3 | | 15.93 | **0.18** |
| Af_6 | Rep1 | 0.64 | 24.40 | **4.14** | Af_6 | Rep1 | 8 | 21.75 | **0.41** |
| | Rep2 | | 22.97 | **3.07** | | Rep2 | | 21.46 | **3.34** |
| | Rep3 | | 24.65 | **7.54** | | Rep3 | | 19.75 | **4.60** |
| Af_6 | Rep1 | 1.61 | 21.08 | **12.52** | Af_6 | Rep1 | 12 | 21.13 | **5.16** |
| | Rep2 | | 21.28 | **13.12** | | Rep2 | | 23.23 | **3.44** |
| | Rep3 | | 21.59 | **13.63** | | Rep3 | | 22.41 | **6.41** |
| Af_6 | Rep1 | 50 | 20.93 | **12.85** | Af_6 | Rep1 | 20 | 23.74 | **3.25** |
| | Rep2 | | 21.51 | **17.41** | | Rep2 | | 23.42 | **1.40** |
| | Rep3 | | 22.33 | **16.09** | | Rep3 | | 22.39 | **4.38** |
| Af_8 | Rep1 | 0.032 | **-1.72** | 0.01 | Af_8 | Rep1 | 1 | **8.67** | 10.41 |
| | Rep2 | | **0.48** | 1.80 | | Rep2 | | **13.37** | NA |
| | Rep3 | | **4.80** | 6.08 | | Rep3 | | 10.39 | **6.56** |
| Af_8 | Rep1 | 0.081 | 5.68 | **-8.78** | Af_8 | Rep1 | 2 | 10.43 | **6.59** |
| | Rep2 | | 13.62 | **-2.82** | | Rep2 | | 15.79 | **7.06** |
| | Rep3 | | 8.74 | **-5.39** | | Rep3 | | 10.01 | **8.64** |
| Af_8 | Rep1 | 0.32 | 5.32 | **-10.92** | Af_8 | Rep1 | 4 | 14.99 | **13.85** |
| | Rep2 | | 6.21 | **-2.45** | | Rep2 | | 19.98 | **15.09** |
| | Rep3 | | 3.10 | **-9.61** | | Rep3 | | 17.31 | **10.18** |
| Af_8 | Rep1 | 0.64 | 14.41 | **-8.39** | Af_8 | Rep1 | 8 | 20.91 | **13.24** |
| | Rep2 | | 14.71 | **4.29** | | Rep2 | | 21.71 | **13.60** |
| | Rep3 | | 16.10 | **10.41** | | Rep3 | | 21.60 | **16.09** |
| Af_8 | Rep1 | 1.61 | 16.11 | **11.80** | Af_8 | Rep1 | 12 | 22.35 | **13.80** |
| | Rep2 | | 16.13 | **10.98** | | Rep2 | | 24.44 | **12.97** |
| | Rep3 | | 13.72 | **13.00** | | Rep3 | | 23.02 | **16.91** |
| Af_8 | Rep1 | 50 | 18.96 | **12.23** | Af_8 | Rep1 | 20 | 23.95 | **14.30** |
| | Rep2 | | 17.68 | **14.51** | | Rep2 | | 24.74 | **15.47** |
| | Rep3 | | 16.37 | **13.84** | | Rep3 | | 23.77 | **11.75** |
| Af_15 | Rep1 | 0.032 | 16.74 | **8.73** | Af_15 | Rep1 | 1 | **17.89** | 19.13 |
| | Rep2 | | 10.57 | **1.15** | | Rep2 | | **19.64** | 20.90 |
| | Rep3 | | 14.33 | **6.95** | | Rep3 | | **20.95** | NA |
| Af_15 | Rep1 | 0.081 | -1.52 | **-3.28** | Af_15 | Rep1 | 2 | **13.47** | NA |
| | Rep2 | | -3.01 | **-10.15** | | Rep2 | | 11.55 | **11.45** |
| | Rep3 | | 8.86 | **1.62** | | Rep3 | | 19.66 | **16.55** |
| Af_15 | Rep1 | 0.32 | 24.02 | **-9.74** | Af_15 | Rep1 | 4 | 19.18 | 12.84 |
| | Rep2 | | 24.03 | **-2.71** | | Rep2 | | 15.32 | 8.01 |
| | Rep3 | | 22.08 | **2.72** | | Rep3 | | **19.76** | NA |
| Af_15 | Rep1 | 0.64 | 27.90 | **-0.55** | Af_15 | Rep1 | 8 | 21.28 | **7.46** |
| | Rep2 | | 27.32 | **6.68** | | Rep2 | | 22.89 | **9.12** |
| | Rep3 | | 29.62 | **9.09** | | Rep3 | | 21.82 | **5.76** |
| Af_15 | Rep1 | 1.61 | 23.67 | **11.51** | Af_15 | Rep1 | 12 | 22.17 | **9.28** |

| | | | | | | | | | |
|---|---|---|---|---|---|---|---|---|---|
| | Rep2 | | 23.18 | **14.77** | | Rep2 | | 23.87 | **10.43** |
| | Rep3 | | 22.59 | **17.92** | | Rep3 | | 21.90 | **10.35** |
| Af_15 | Rep1 | 50 | 24.46 | **10.89** | Af_15 | Rep1 | 20 | 23.97 | **4.93** |
| | Rep2 | | 23.89 | **11.60** | | Rep2 | | 22.36 | **10.51** |
| | Rep3 | | 23.05 | **18.03** | | Rep3 | | 24.85 | **9.63** |
| Af_3 | Rep1 | 0.032 | 1.12 | **-7.74** | Af_3 | Rep1 | 1 | 2.92 | **-20.54** |
| | Rep2 | | 9.50 | **1.97** | | Rep2 | | -0.03 | **-2.78** |
| | Rep3 | | -1.99 | **-14.76** | | Rep3 | | **4.37** | 5.58 |
| Af_3 | Rep1 | 0.081 | 7.32 | **-6.55** | Af_3 | Rep1 | 2 | -3.26 | **-12.62** |
| | Rep2 | | 8.91 | **0.57** | | Rep2 | | **3.89** | 5.36 |
| | Rep3 | | 6.49 | **6.01** | | Rep3 | | 1.71 | **-4.18** |
| Af_3 | Rep1 | 0.32 | 27.14 | **-6.82** | Af_3 | Rep1 | 4 | 12.15 | **-17.98** |
| | Rep2 | | 24.53 | **-6.38** | | Rep2 | | 13.35 | **-7.00** |
| | Rep3 | | 22.36 | **-4.01** | | Rep3 | | 17.51 | **-14.83** |
| Af_3 | Rep1 | 0.64 | 28.00 | **6.56** | Af_3 | Rep1 | 8 | 19.69 | **4.41** |
| | Rep2 | | 28.34 | **4.75** | | Rep2 | | 21.51 | **5.16** |
| | Rep3 | | 27.94 | **11.57** | | Rep3 | | 21.30 | **6.73** |
| Af_3 | Rep1 | 1.61 | 19.84 | **3.80** | Af_3 | Rep1 | 12 | 24.14 | **-0.45** |
| | Rep2 | | 20.96 | **4.35** | | Rep2 | | 23.29 | **1.83** |
| | Rep3 | | 17.19 | **8.95** | | Rep3 | | 23.11 | **3.40** |
| Af_3 | Rep1 | 50 | 19.58 | **7.09** | Af_3 | Rep1 | 20 | 24.11 | **-2.06** |
| | Rep2 | | 18.56 | **6.58** | | Rep2 | | 23.82 | **1.88** |
| | Rep3 | | 18.67 | **9.70** | | Rep3 | | 24.41 | **3.77** |
| Af_12 | Rep1 | 0.032 | 5.47 | **4.47** | Af_12 | Rep1 | 1 | -4.00 | **-4.29** |
| | Rep2 | | -0.13 | **-3.07** | | Rep2 | | 1.95 | **-1.54** |
| | Rep3 | | 2.53 | **1.72** | | Rep3 | | **4.52** | 5.69 |
| Af_12 | Rep1 | 0.081 | 6.98 | **2.24** | Af_12 | Rep1 | 2 | 9.86 | **3.58** |
| | Rep2 | | -4.65 | **-7.94** | | Rep2 | | -1.29 | **-6.40** |
| | Rep3 | | 12.06 | **0.69** | | Rep3 | | 4.43 | **-7.64** |
| Af_12 | Rep1 | 0.32 | 21.86 | **-9.55** | Af_12 | Rep1 | 4 | 16.46 | **-13.26** |
| | Rep2 | | 21.62 | **-7.25** | | Rep2 | | 15.09 | **-10.35** |
| | Rep3 | | 21.30 | **-4.00** | | Rep3 | | 15.28 | **-5.58** |
| Af_12 | Rep1 | 0.64 | 27.69 | **10.63** | Af_12 | Rep1 | 8 | 21.96 | **-4.99** |
| | Rep2 | | 26.79 | **9.05** | | Rep2 | | 19.45 | **-3.18** |
| | Rep3 | | 27.28 | **17.83** | | Rep3 | | 20.02 | **-7.04** |
| Af_12 | Rep1 | 1.61 | 21.32 | **9.08** | Af_12 | Rep1 | 12 | 20.34 | **-1.19** |
| | Rep2 | | 22.22 | **10.66** | | Rep2 | | 20.67 | **-4.94** |
| | Rep3 | | 18.22 | **10.19** | | Rep3 | | 22.66 | **-3.55** |
| Af_12 | Rep1 | 50 | 21.35 | **11.13** | Af_12 | Rep1 | 20 | 23.17 | **-3.64** |
| | Rep2 | | 20.10 | **10.58** | | Rep2 | | 22.28 | **-6.11** |
| | Rep3 | | 20.24 | **14.81** | | Rep3 | | 23.30 | **-5.63** |
| Af_18 | Rep1 | 0.032 | 14.64 | **6.49** | Af_18 | Rep1 | 1 | -4.06 | **-5.44** |
| | Rep2 | | 17.11 | **8.37** | | Rep2 | | **5.13** | NA |
| | Rep3 | | 14.44 | **10.55** | | Rep3 | | **-1.28** | -0.85 |
| Af_18 | Rep1 | 0.081 | 19.33 | **7.66** | Af_18 | Rep1 | 2 | 6.60 | **6.00** |
| | Rep2 | | 15.90 | **1.94** | | Rep2 | | 1.93 | **-11.40** |
| | Rep3 | | 17.79 | **-4.98** | | Rep3 | | 7.75 | **5.02** |
| Af_18 | Rep1 | 0.32 | 28.66 | **0.24** | Af_18 | Rep1 | 4 | 13.55 | **-18.70** |
| | Rep2 | | 27.97 | **-7.03** | | Rep2 | | 17.24 | **-5.68** |
| | Rep3 | | 26.51 | **-1.76** | | Rep3 | | 13.99 | **-8.64** |
| Af_18 | Rep1 | 0.64 | 30.08 | **10.35** | Af_18 | Rep1 | 8 | 19.40 | **-4.91** |
| | Rep2 | | 28.94 | **13.79** | | Rep2 | | 18.92 | **2.04** |
| | Rep3 | | 28.60 | **17.24** | | Rep3 | | 19.51 | **-1.02** |
| Af_18 | Rep1 | 1.61 | 21.95 | **12.91** | Af_18 | Rep1 | 12 | 21.34 | **-3.73** |
| | Rep2 | | 22.11 | **17.57** | | Rep2 | | 20.83 | **-3.81** |
| | Rep3 | | 22.98 | **16.66** | | Rep3 | | 19.48 | **-0.67** |
| Af_18 | Rep1 | 50 | 21.56 | **11.86** | Af_18 | Rep1 | 20 | 23.20 | **-5.36** |
| | Rep2 | | 21.95 | **15.04** | | Rep2 | | 22.01 | **-3.94** |
| | Rep3 | | 21.08 | **15.97** | | Rep3 | | 22.63 | **1.78** |

42 **c)**

| **Temperature experiment – Growth rates** | | | |
|---|---|---|---|
| **Isolate** | **Temperature [°C]** | **AIC linear model** | **AIC logistic model** |
| Af_5 | 5 | **14.50** | 14.95 |
| | 10 | 38.62 | **23.31** |
| | 15 | 42.30 | **16.10** |
| | 20 | 36.98 | **11.30** |
| | 22.5 | 35.81 | **9.44** |
| Af_8 | 5 | 14.50 | **9.27** |
| | 10 | 43.21 | **20.07** |
| | 15 | 41.76 | **10.08** |
| | 20 | 36.90 | **14.12** |
| | 22.5 | 35.71 | **9.70** |
| Af_15 | 5 | **8.06** | 9.17 |
| | 10 | 37.51 | **19.89** |
| | 15 | 35.42 | **7.07** |
| | 20 | 33.34 | **13.90** |
| | 22.5 | 32.11 | **3.52** |
| Af_3 | 5 | 14.09 | **4.97** |
| | 10 | 37.12 | **16.75** |
| | 15 | 36.48 | **7.69** |
| | 20 | 33.27 | **9.93** |
| | 22.5 | 31.46 | **9.56** |
| Af_18 | 5 | 18.15 | **15.88** |
| | 10 | 42.45 | **21.72** |
| | 15 | 40.69 | **5.70** |
| | 20 | 37.34 | **8.61** |
| | 22.5 | 34.42 | **9.47** |

| **Temperature experiment – TPC** | | | | | |
|---|---|---|---|---|---|
| **Isolate** | Af_5 | Af_8 | Af_15 | Af_3 | Af_18 |
| **AIC 1** | 12.82 | 14.36 | 12.84 | 12.32 | 14.08 |
| **AIC 2** | 10.9 | 12.66 | 11.25 | 10.29 | 12.11 |
| **AIC 3** | 9.83 | 11.28 | 10.03 | 9.27 | 11.09 |
| **AIC 4** | 14.69 | 16.28 | 15.2 | 13.66 | 16.11 |
| **AIC 5** | 14.1 | 15.82 | 16.36 | 13.31 | 15.96 |
| **AIC 6** | 8.46 | 9.46 | 8.12 | 7.45 | 9.59 |
| **AIC 7** | 9.9 | 11.37 | 9.81 | 9.45 | 11.16 |
| **AIC 8** | 3.75 | 12.86 | 1.84 | 2.29 | 4.75 |
| **AIC 9** | 8.79 | 9.62 | 8.44 | 7.87 | 10 |
| **AIC 10** | -4.06 | -4.88 | 1.83 | -8.19 | -0.66 |
| **AIC 11** | -21.23 | -13.48 | -26.34 | -28.39 | -16.17 |
| **AIC 12** | 14.07 | 15.56 | 14.31 | 13.54 | 15.34 |
| **AIC 13** | 11.96 | 12.88 | 24.18 | 14.41 | 16.25 |
| **AIC 14** | 12.1 | 13.78 | 13.16 | 11.41 | 13.52 |
| **AIC 15** | 11.32 | 12.4 | 12.06 | 10.79 | 12.84 |
| **AIC 16** | -4.06 | -4.88 | 1.83 | -8.19 | -0.66 |

**Table S2.** Sites under selection involved in defense/immune responses between subpopulations. The last five rows report the additional sites under selection found between subpopulations 1 and 3. Selection: positive (+), balancing (0). For mutations, the AA substitutions are reported.

| Contig | Gene | Function | Annotation derived from | Subpopulations, Selection | Mutation |
|---|---|---|---|---|---|
| NKIB01012762.1 1410 | **BAM3**: Leucine-rich repeat receptor-like serine/threonine-protein kinase BAM3 | Role in plant defense to fungal exposure | *Arabidopsis thaliana* | Subp1 - Subp3: +; Subp2 - Subp4: + | Synonymous and nonsynonymous (from E to D) |
| NKIB01009463.1 9326 | **MIK2**: MDIS1-interacting receptor like kinase 2 | Crucial component of early immune responses to a fungal-derived elicitor | *Arabidopsis thaliana* | Subp1 - Subp3: +; Subp1 – Subp4: +; Subp2 - Subp4: + | Nonsynonymous (from D to E; from S to P) |
| NKIB01006846.1 12531 | **NLRC3:** NLR family CARD domain-containing protein 3 | Encodes a NOD-like receptor family member. The encoded protein is a cytosolic regulator of innate immunity | *Homo sapiens* | Subp1 - Subp3: +; Subp2 – Subp3: +; Subp3 - Subp4: + | Nonsynonymous (from D to E) |
| NKIB01007086.1 7984 | **NLRC3:** NLR family CARD domain-containing protein 3 | Encodes a NOD-like receptor family member. The encoded protein is a cytosolic regulator of innate immunity | *Homo sapiens* | Subp2 - Subp4: + | Nonsynonymous (from E to G) |
| NKIB01008260.1 8281 | **NLRC3:** NLR family CARD domain-containing protein 3 | Encodes a NOD-like receptor family member. The encoded protein is a cytosolic regulator of innate immunity | *Homo sapiens* | Subp3 - Subp4: 0 | Nonsynonymous (from D to E) |
| NKIB01013721.1 2824 | **NLRC3:** NLR family CARD domain-containing protein 3 | Encodes a NOD-like receptor family member. The encoded protein is a cytosolic regulator of innate immunity | *Homo sapiens* | Subp1 – Subp2: 0; Subp1 - Subp3: 0; Subp1 - Subp4: 0; Subp2 – Subp3: 0; Subp2 - Subp4: 0 | Nonsynonymous (from R to Q) |
| NKIB01012421.1 991 | **FLS2:** LRR receptor-like serine/threonine-protein kinase FLS2 | Pathogen-associated molecular pattern | *Arabidopsis thaliana* | Subp1 - Subp2: +; Subp2 - Subp3: +; Subp2 – Subp4: +; Subp3 - Subp4: + | Nonsynonymous (from K to Q; from K to *) |
| NKIB01012262.1 4723, 4724, 4725, 4728 | **NPR4:** Ankyrin repeat-containing protein NPR4 | Involved in plant immunity. NPR4 is required for basal defense against pathogens | *Oryza sativa subsp. japonica* | Subp1 - Subp2: NA; Subp1 – Subp4: 0; Subp2 – Subp3: +; Subp2 - Subp4: +; Subp3 - Subp4: + | Synonymous and nonsynonymous (from S to L; from S to T; from T to P) |
| NKIB01002010.1 1291 | **At5g63930:** Probable leucine-rich repeat receptor-like protein kinase | Involved in defense response | *Arabidopsis thaliana* | Subp2 - Subp4: + | Nonsynonymous (from V to F) |
| NKIB01011144.1 2146, 4634 | **AT4G04500.1 (CRK37):** Cysteine-rich receptor-like protein kinase 37 | Regulate several biological processes, including plant development and stress adaptation | *Arabidopsis thaliana* | Subp1 - Subp3: 0 | |
| NKIB01000724.1 255 | **FIB1:** rRNA 2'-O-methyltransferase fibrillarin 1 | RNA methylation | *Arabidopsis thaliana* | Subp1 - Subp3: + | |
| NKIB01009964.1 7847 | **PCCB:** Propionyl-CoA carboxylase beta chain%2C mitochondrial | Involved in the degradation of several essential amino acids (threonine, valine, isoleucine, and methionine), fatty acids of odd numbered chain lengths, and cholesterol | *Sus scrofa* | Subp1 - Subp3: + | |
| NKIB01012394.1 7565 | **FKBP19:** Peptidyl-prolyl cis-trans isomerase FKBP19%2C chloroplastic | Protein synthesis, folding, and degradation | *Arabidopsis thaliana* | Subp1 - Subp3: 0 | |
| NKIB01013520.1 41753 | **AT1G22620.1 (SAC1):** Phosphatidylinositol-3-phosphatase SAC1 | Involved in cell morphogenesis, cell wall synthesis, and actin organization | *Arabidopsis thaliana* | Subp1 - Subp3: + | |

**Table S3.** Thermal coefficients for some of the isolates obtained using the R packages *rTPC* and *nls.multstart* [2]. Thermal performance breadth is the range of temperatures over which the rate of a curve is at least 0.8 of the peak.

| Isolate | Thermal optimum (Topt) [°C] | Critical thermal maximum (Tmax) [°C] | Thermal performance breadth |
|---|---|---|---|
| Af_5 | 21.70 | 24.85 | 12.02 |
| Af_8 | 21.74 | 24.82 | 9.60 |
| Af_15 | 21.26 | 24.70 | 11.75 |
| Af_3 | 21.66 | 24.76 | 10.97 |
| Af_18 | 21.77 | 24.81 | 9.40 |
